# Clustering by phenotype and genome-wide association study in autism

**DOI:** 10.1101/614958

**Authors:** Akira Narita, Masato Nagai, Satoshi Mizuno, Soichi Ogishima, Gen Tamiya, Masao Ueki, Rieko Sakurai, Satoshi Makino, Taku Obara, Mami Ishikuro, Chizuru Yamanaka, Hiroko Matsubara, Yasutaka Kuniyoshi, Keiko Murakami, Fumihiko Ueno, Aoi Noda, Tomoko Kobayashi, Mika Kobayashi, Takuma Usuzaki, Hisashi Ohseto, Atsushi Hozawa, Masahiro Kikuya, Hirohito Metoki, Shigeo Kure, Shinichi Kuriyama

## Abstract

**Background:** Autism spectrum disorder (ASD) has phenotypically and genetically heterogeneous characteristics. A simulation study demonstrated that attempts to categorize patients with a complex disease into more homogeneous subgroups could have more power to elucidate hidden heritability.

**Methods:** We conducted cluster analyses using the k-means algorithm with a cluster number of 15 based on phenotypic variables from the Simons Simplex Collection (SSC). As a preliminary study, we conducted a conventional genome-wide association study (GWAS) with a dataset of 597 ASD cases and 370 controls. In the second step, we divided cases based on the clustering results and conducted GWAS in each of the subgroups vs controls (cluster-based GWAS). We also conducted cluster-based GWAS on another SSC dataset of 712 probands and 354 controls in the replication stage.

**Results:** In the preliminary study, we observed no significant associations. In the second step of cluster-based GWASs, we identified 65 chromosomal loci, which included 30 intragenic loci located in 21 genes and 35 intergenic loci that satisfied the threshold of *P*<5.0×10^−8^. Some of these loci were located within or near previously reported candidate genes for ASD: *CDH5, CNTN5, CNTNAP5, DNAH17, DPP10, DSCAM, FOXK1, GABBR2, GRIN2A*5, *ITPR1, NTM, SDK1, SNCA* and *SRRM4.* Of these 65 significant chromosomal loci, rs11064685 located within the *SRRM4* gene had a significantly different distribution in the cases vs. controls in the replication cohort.

**Conclusions:** These findings suggest that clustering may successfully identify subgroups with relatively homogeneous disease etiologies. Further cluster validation and replication studies are warranted in larger cohorts.

## Introduction

Autism spectrum disorder (ASD) has heterogeneous characteristics in terms of both phenotypic features and genetics. ASD is mainly characterized by difficulties in communication and repetitive behaviors (1), but ASD also shows many other symptoms (2). Regarding genetics, previous studies have not consistently identified genetic variants that are associated with an increased risk of ASD (3), although several lines of evidence suggest that genetic factors strongly contribute to the increased risk of ASD. Monozygotic twins have higher concordance rates of ASD (92%) than dizygotic twins (10%) (4). The recurrence risk ratio is 22 for ASD among siblings (5). The Human Gene module of the Simons Foundation Autism Research Initiative (SFARI) Gene provides a comprehensive reference for suggested human ASD-related genes in an up-to-date manner (6) and currently demonstrates ∼1,000 genes that may have links to ASD, potentially indicating the heterogeneity of ASD. In addition to phenotype and genotype heterogeneities, ASD shows heterogeneous responses to interventions. Several kinds of pharmacological treatments are suggested, but the effects of these treatments are controversial (7).

If the heterogeneous phenotypes and responses to treatment in some way correspond to differences in genotype, grouping persons with ASD according to phenotype and responses to treatment variables may increase the chances of identifying genetic susceptibility factors. Traylor and colleagues (8) demonstrated that attempts to categorize patients with a complex disease into more homogeneous subgroups could have more power to elucidate the hidden heritability in a simulation study. Several studies on Alzheimer’s disease, neuroticism, or asthma indicated that items or symptoms were to some degree more useful for identifying high-impact genetic factors than broadly defined diagnoses (9-11), although a study of ASD demonstrated modest effects of two-way stratification by individual symptoms (12). Additionally, medical researchers have begun to use machine learning methods (13), which is an artificial intelligence technique that can reveal masked patterns of data sets. In view of the abovementioned circumstances, clustering algorithms of machine learning could be hypothesized to make novel and more genetically homogeneous clusters, but these algorithms using phenotypic variables have not, to the best of our knowledge, been applied to subgrouping ASD to date.

We therefore explored whether grouping persons with ASD using a clustering algorithm with phenotype and responses to treatment variables can be used to discriminate more genetically homogeneous persons with ASD. In the present study, we conducted cluster-based genome-wide association studies (named cluster-based GWASs) using real data based on the concept of a previous simulation study (8) adopting a machine learning k-means (14) algorithm for cluster analysis.

## Methods and Materials

We conducted the present study in accordance with the guidelines of the Declaration of Helsinki (15) and all other applicable guidelines. The protocol was reviewed and approved by the institutional review board of Tohoku University Graduate School of Medicine, and written informed consent was obtained from all participants over the age of 18 by the Simons Foundation Autism Research Initiative (SFARI) (16). For participants under the age of 18, informed consent was obtained from a parent and/or legal guardian. Additionally, for participants 10 to 17 years of age, informed assent was obtained from the individuals.

### Datasets

We used phenotypic variables, history of treatment, and genotypic data from the Simons Simplex Collection (SSC) (16). The SSC establishes a repository of phenotypic data and genetic data/samples from mainly simplex families.

The SSC data were publicly released in October 2007 and are directly available from the SFARI. From the SSC dataset, we used data from 614 affected white male probands who had no missing information regarding Autism Diagnostic Interview-Revised (ADI-R) scores (17) and vitamin treatment (18, 19) and 391 unaffected brothers for whom genotype data, generated by the Illumina Human Omni2.5 (Omni2.5) array, were available for subsequent clustering and genetic analyses. We excluded participants whose ancestries were estimated to be different from the other participants using principal component analyses (PCAs) performed by EIGENSOFT version 7.2.1 (20, 21) for the genotype data. Based on the PCAs, we excluded data beyond 4 standard deviations of principal components 1 or 2 (Figure S1). Therefore, we used data from 597 probands and 370 unaffected brothers.

In the replication study, we used another SSC dataset genotyped using the Illumina 1Mv3 (1Mv3) array. In the dataset, data from 735 affected male probands with no missing information regarding ADI-R scores or vitamin treatment and 387 unaffected brothers were available. After conducting PCA, we excluded data beyond 4 standard deviations of principal components 1 or 2 as outliers. In this way, we used data from 712 probands and 354 unaffected brothers in the replication study.

### Clustering

We conducted cluster analyses using phenotypic variables of ADI-R (17) scores and history of vitamin treatment (18, 19). We chose these variables because the ADI-R is one of the most reliable estimates of ASD and has the ability to evaluate substructure domains of ASD (17). Among the ADI-R scores, “the total score for the Verbal Communication Domain of the ADI-R minus the total score for the Nonverbal Communication Domain of the ADI-R”, “the total score for the Nonverbal Communication Domain of the ADI-R”, “the total score for the Restricted, Repetitive, and Stereotyped Patterns of Behavior Domain of the ADI-R”, and “the total score for the Reciprocal Social Interaction Domain of the ADI-R” were included in the preprocessed dataset.

Among the treatments, we selected the variable of history of vitamin treatment because we recently found that a cluster of persons with ASD is associated with potential responsiveness to vitamin B6 treatment (18, 19). The history of treatment is not always compatible with responsiveness, but we considered that continuous treatment indicates responsiveness to some degree. The SSC dataset includes history of treatment but not variables of responsiveness.

We applied the machine learning k-means (14) algorithm to conduct a cluster analysis to divide the dataset including data from ASD persons into subgroups using phenotypic variables and history of treatment. The k-means algorithm requires cluster numbers determined by researchers. When using k-means algorithms, we set a priori the cluster numbers of 2, 3, 4, 5, 10, 15, and 20. We performed the analyses using the scikit-learn toolkit in Python 2.7 (Supplementary Information S1) (22).

Clustering is an exploratory data analysis technique, and the validity of the clustering results may be judged by external knowledge, such as the purpose of the segmentation (23). Several methods have proposed to prespecify a cluster number (k), such as visual examination of the data, and likelihood and error-based approaches; however, these methods do not necessarily provide results that are consistent with each other (24). Although there are measures for evaluating the quality of the clusters (25), the number of clusters should also be determined according to the research purposes. We regarded the inflation factor (λ) of quantile-quantile (Q-Q) plots of the logarithm of the P-value to base 10 (-log10P) as one of the indicators of successful clustering in the present study. We calculated λ for each cluster number.

When conducting clustering, we combined the two datasets of male probands, one genotyped using the Omni2.5 array and the other genotyped using the 1Mv3 array. After clustering, we redivided the new dataset according to the SNP arrays used. In the discovery stage, we used the Omni2.5 dataset and the 1Mv3 dataset in the replication stage.

### Genotype data and quality control

We used the SSC dataset, in which probands and unaffected brothers had already been genotyped in other previous studies (16, 26). In the discovery stage, we used the dataset genotyped by the Omni2.5 array, which has 2,383,385 probes. We excluded SNPs with a minor allele frequency (MAF) < 0.01, call rate < 0.95, and Hardy-Weinberg equilibrium test P < 0.000001.

In the replication study, where we used the dataset genotyped using the 1Mv3 array, we applied the same cutoff values for quality control as those used in the discovery stage. The 1Mv3 array includes 1,147,689 SNPs. The Omni2.5 array and the 1Mv3 array shared 675,923 SNPs.

### Statistical analysis

As a preliminary study, we conducted a conventional GWAS in the whole Omni2.5 dataset, with a total of 597 male probands and 370 unaffected brothers. Here, we used the brothers of the cases as controls, in contrast to many previous studies in which genetically unrelated controls were used. We thus adopted the sib transmission disequilibrium test (sib-TDT) (27), a family-based association test, to take into account familial relationships among the participants. In the second step, in the discovery stage, we conducted cluster-based GWAS in each subgroup of the cases, which had been divided using the k-means (14) algorithm, and the controls. As mentioned above, the controls were the brothers of the cases, and we then excluded the unaffected brothers of the cases belonging to the subgroup being analyzed. Details of the study design are shown in Figure 1. We applied the Cochran-Armitage trend test (28), which examines the risk of disease in those who do not have the allele of interest, those who have a single copy, and those who are homozygous.

**Figure 1.**
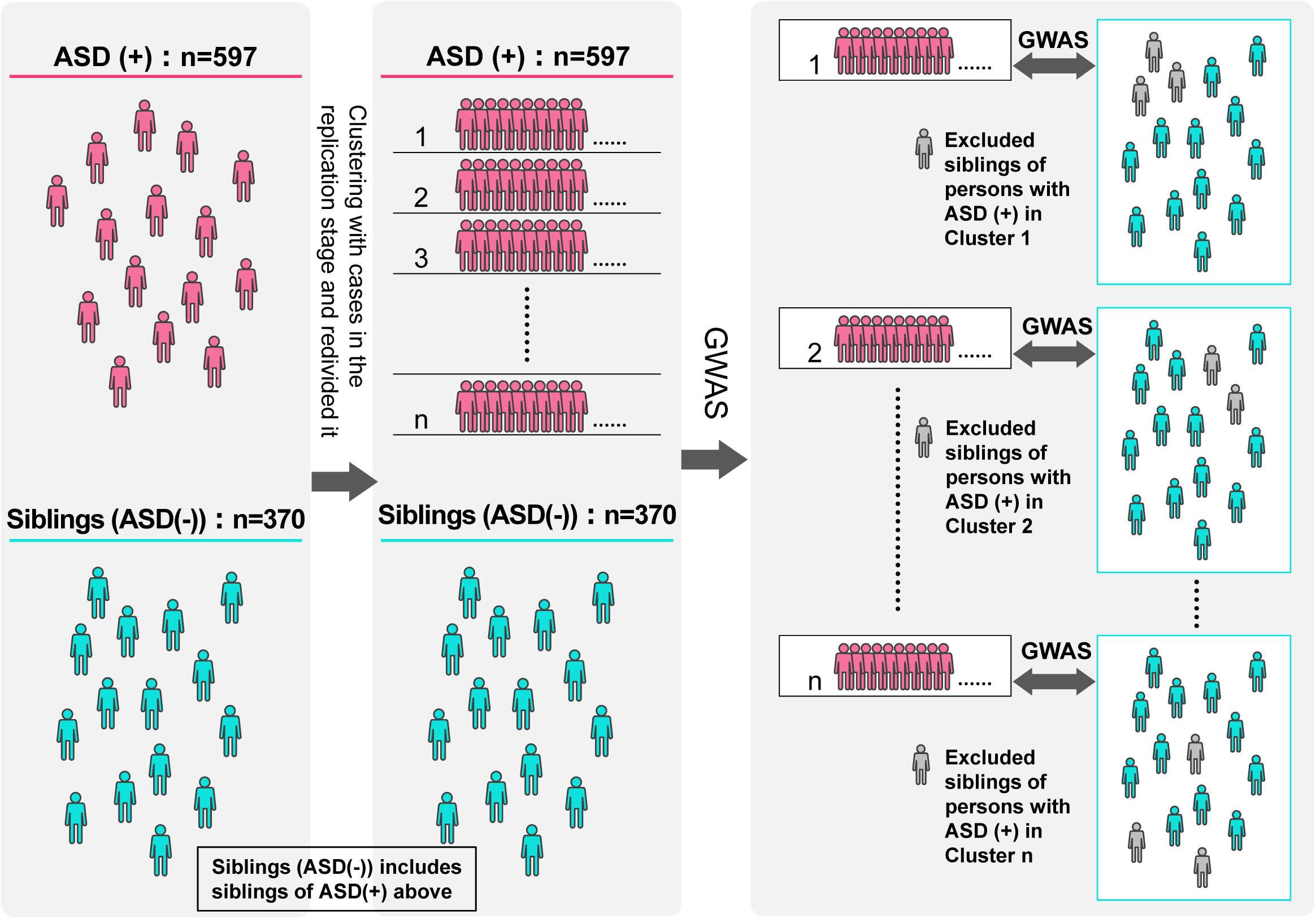
Details of the cluster-based GWAS in the present discovery stage in the Simons Simplex Collection dataset. In the present study, a GWAS using each subgroup of the probands vs the unaffected brothers as controls without the brothers of the members of the subgroup was designated a “cluster-based GWAS”. This panel shows the detailed methods of the cluster-based GWAS in the discovery stage.

We further tested the significantly associated loci found in the discovery studies in the replication stage. The level of significance for association was set as P <0.05 in the replication studies.

Association analyses were performed with the PLINK software package (29). The detected SNPs were subsequently annotated using ANNOVAR (30). Manhattan plots and Q-Q plots were generated using the ‘qqman’ package in R version 3.0.2.

### Data availability

All the data used in the study are available only to those granted access by the Simons Foundation.

## Results

### Cluster-based genome-wide association study

As a preliminary study, we conducted a conventional GWAS with the Omni2.5 dataset using the sib-TDT. We observed no significant associations (Figure 2). Although we adopted the sib-TDT here because we used the brothers of the cases as controls, we also used the Cochran-Armitage trend test and found that the -log10P values were distributed downward compared with the expected values, as shown in Figure S2.

**Figure 2.**
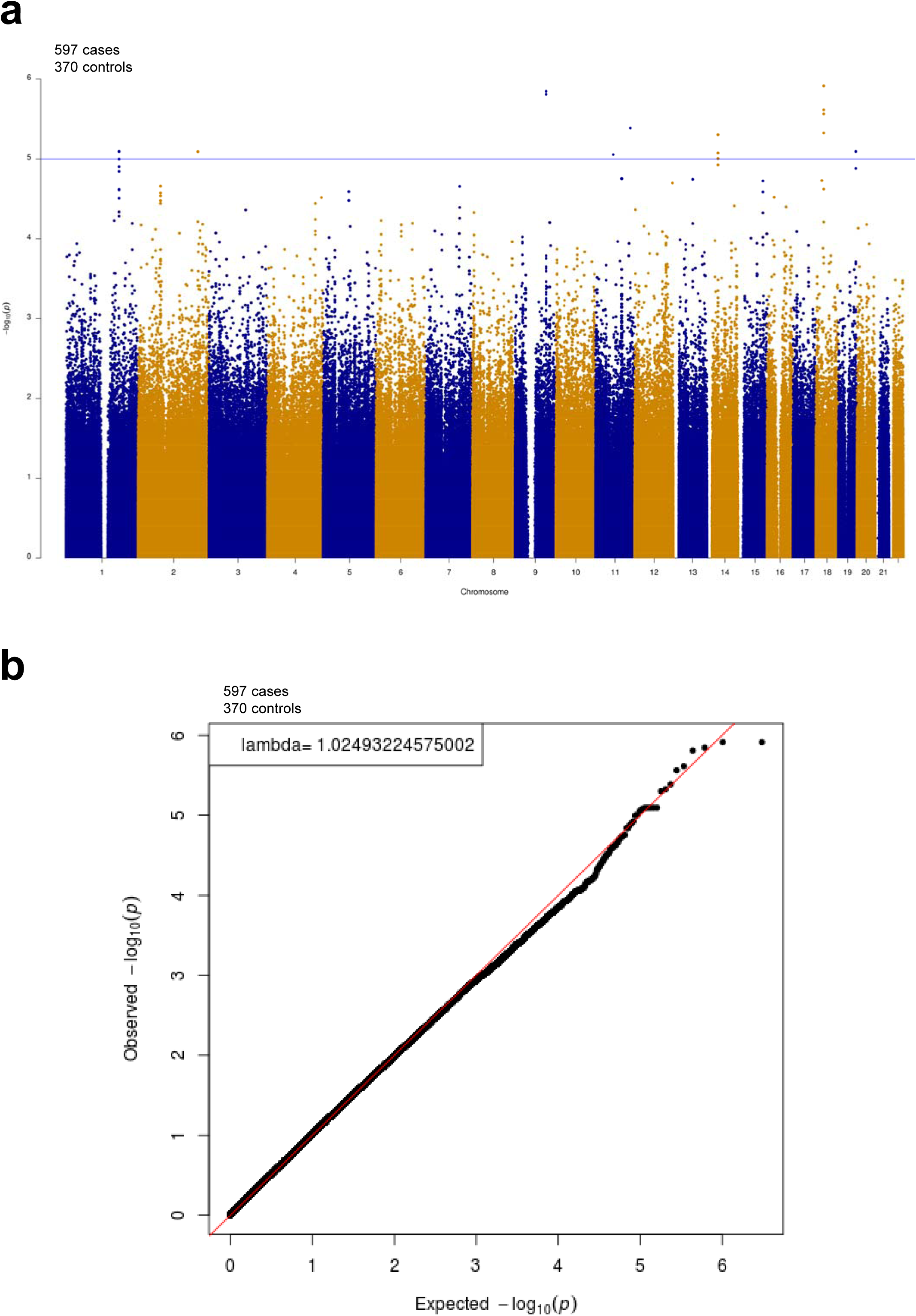
Manhattan plots (a) and corresponding quantile-quantile plots (b) in GWAS for all male probands vs their unaffected brothers using the sib transmission/disequilibrium test. We conducted a GWAS in the Simons Simplex Collection dataset of 597 male probands and 370 unaffected brothers genotyped by the Illumina Human Omni2.5 array using the sib transmission/disequilibrium test (sib-TDT). We observed no significant associations in this GWAS with the genome-wide threshold of P < 5.0 × 10^−8^.

We also applied the sib-TDT to cluster 1, which was obtained by dividing all the cases using k-means with k of 15, and all the controls and found that the observed -logP values were lower than expected, as shown in Figure S2. Since the sib-TDT may efficiently work in a population consisting of a substantial number of sibs, a limited number of brothers of the probands among all the controls probably contributed to a substantial loss of power. Thus, we excluded the brothers of the probands in each subset from the controls so that each subset of probands has no genetic relations with the rest of the controls and conducted the Cochran-Armitage trend test, as in many other studies. In the present study, therefore, we applied the sib-TDT to the GWAS of the whole dataset, whereas in the cluster-based GWAS, we excluded in turn the unaffected brothers of the cases belonging to the subgroup being analyzed and used the Cochran-Armitage trend test to account for the relationships between participants.

Under the hypothesis that ASD consists of hundreds of subgroups (16), we compared λ values giving larger numbers of clusters as priority. The λ for the cluster-based GWAS with a k of 20 ranged from 1.015 to 1.107, and the average was 1.053 (Table 1), indicating that the rate of false positives was relatively high. Several lines of evidence suggest that regarding an appropriate threshold of inflation factor λ, empirically, a value less than 1.050 is deemed safe for avoiding false positives (31-33).

**Table 1.** Number of genome-wide significant loci and λ values according to the number of clusters using the k-means algorithm and the Omni2.5 dataset with MAF <0.01 deleted.

| No. of Clusters | 1 | 2 | 3 | 4 | 5 | 10 | 15 | 20 |
| --- | --- | --- | --- | --- | --- | --- | --- | --- |
| Number of genome-wide significant loci | 0 | 0 | 2 | 5 | 1 | 26 | 65 | 211 |
| Mean $\lambda$ value (min-max) | 1.032 | 1.057<br>(1.052-1.062) | 1.036<br>(1.027-1.045) | 1.035<br>(1.029-1.041) | 1.021<br>(1.012-1.033) | 1.024<br>(1.000-1.042) | 1.038<br>(1.017-1.091) | 1.053<br>(1.015-1.107) |

$\lambda$ values were calculated from the sibling-based transmission disequilibrium test in the number of clusters 1 or the Cochran-Armitage trend test in the number of clusters from 2 to 20.

In contrast, λ with k of 15 ranged from 1.017 to 1.091, and the average was 1.038, which was below 1.050 (Table 1 and Figure 3).

**Figure 3.**
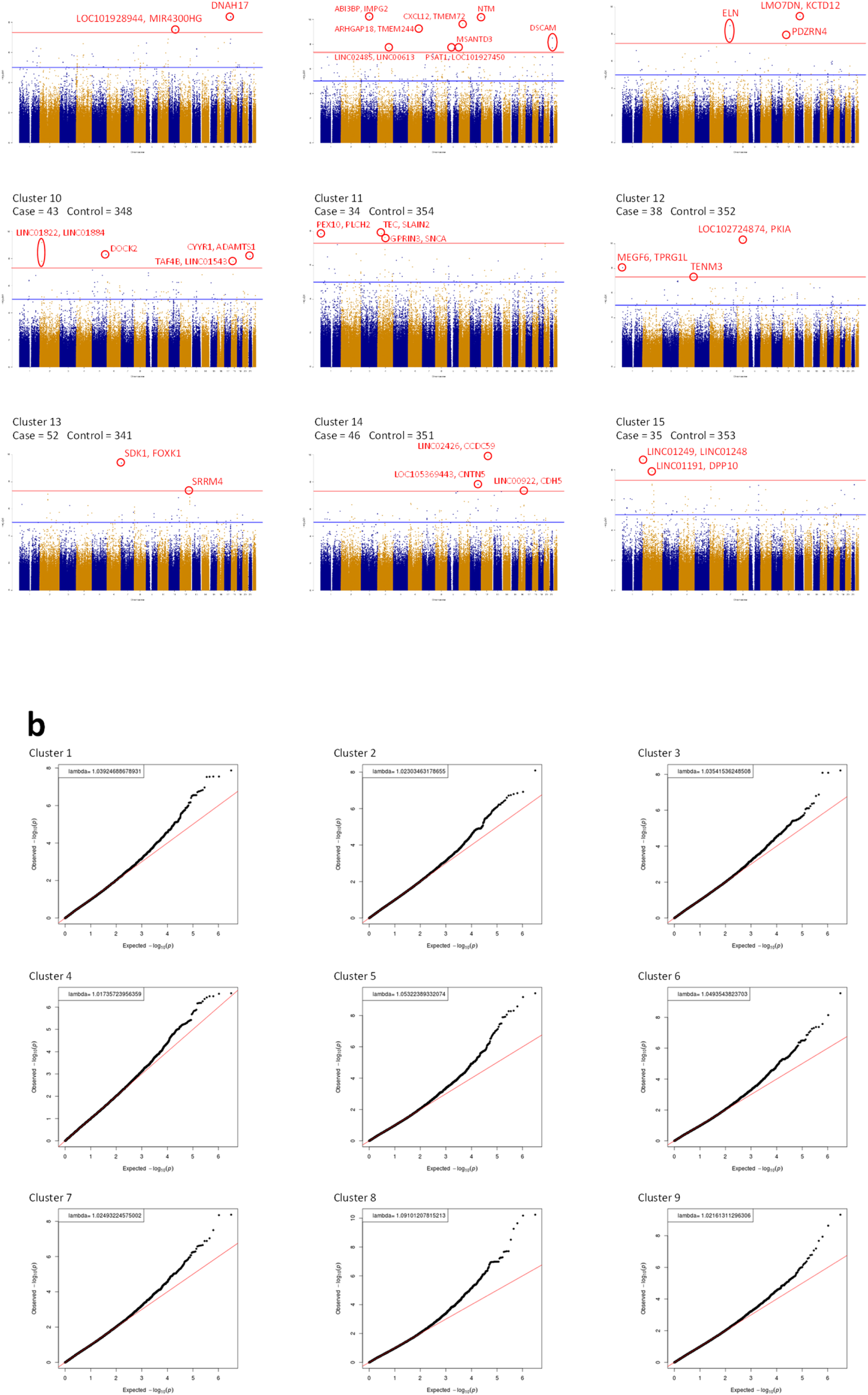
Manhattan plots (a) and corresponding quantile-quantile plots (b) for cluster-based GWASs with a cluster number of 15. We performed cluster analysis using k-means with a cluster number of 15 and conducted cluster-based GWAS. Among 15 clusters, significant associations were observed in 14 clusters. In total, we observed 65 chromosomal loci, labeled in the figure, that satisfied the threshold of P < 5.0 × 10^−8^. The red lines indicate the threshold for genome-wide significance (P < 5.0 × 10^−8^).

According to the above results, we considered the cluster-based GWAS using the Cochran-Armitage trend test, coupled with k-means cluster analysis with k of 15, to be the most appropriate approach to the present dataset. The characteristics of each cluster are presented in Table 2.

**Table 2.** Characteristics of each of 15 k-means clusters in the Omni2.5 dataset.

| Cluster No. | n | Verbal score from ADI-R |  |  |  | Nonverbal score from ADI-R |  |  |  | Restricted and repetitive patterns of behavior score from ADI-R |  |  |  | Social score from ADI-R |  |  |  | Vitamin B6 treatment (%) |
| --- | --- | --- | --- | --- | --- | --- | --- | --- | --- | --- | --- | --- | --- | --- | --- | --- | --- | --- |
|  |  | Mean (SD) | Median (p25-p75) | Min | Max | Mean (SD) | Median (p25-p75) | Min | Max | Mean (SD) | Median (p25-p75) | Min | Max | Mean (SD) | Median (p25-p75) | Min | Max |  |
| All | 597 | 7.7 (2.1) | 8.0 (6.0-9.0) | 0 | 12 | 8.9 (3.3) | 9.0 (6.0-12.0) | 0 | 14 | 6.8 (2.5) | 7.0 (5.0-8.0) | 1 | 12 | 19.8 (5.3) | 20.0 (16.0-24.0) | 8 | 30 | 59.6 |
| 1 | 33 | 7.4 (2.2) | 7.0 (6.0-10.0) | 3 | 11 | 4.4 (1.6) | 4.0 (3.0-6.0) | 1 | 7 | 8.5 (1.6) | 8.0 (7.0-10.0) | 6 | 12 | 14.0 (1.5) | 14.0 (13.0-15.0) | 11 | 17 | 60.6 |
| 2 | 49 | 8.9 (1.3) | 9.0 (8.0-10.0) | 6 | 12 | 12.3 (1.5) | 12.0 (11.0-14.0) | 9 | 14 | 6.2 (1.3) | 6.0 (6.0-7.0) | 3 | 8 | 27.1 (1.3) | 27.0 (26.0-28.0) | 24 | 30 | 79.6 |
| 3 | 45 | 6.0 (1.9) | 6.0 (5.0-7.0) | 2 | 10 | 8.8 (1.5) | 9.0 (8.0-10.0) | 6 | 12 | 5.0 (1.5) | 5.0 (4.0-6.0) | 2 | 7 | 16.8 (1.1) | 17.0 (16.0-18.0) | 15 | 19 | 64.4 |
| 4 | 59 | 9.0 (1.5) | 9.0 (8.0-10.0) | 6 | 12 | 8.1 (1.5) | 8.0 (7.0-9.0) | 4 | 10 | 8.8 (1.9) | 8.0 (8.0-10.0) | 5 | 12 | 23.8 (1.4) | 24.0 (23.0-25.0) | 21 | 27 | 57.6 |
| 5 | 28 | 7.3 (1.1) | 7.0 (6.5-8.0) | 5 | 9 | 9.1 (1.7) | 9.0 (8.0-10.0) | 7 | 13 | 6.1 (2.3) | 6.0 (5.0-7.0) | 1 | 12 | 12.7 (1.7) | 13.0 (12.0-14.0) | 9 | 15 | 60.7 |
| 6 | 29 | 7.7 (1.9) | 8.0 (7.0-9.0) | 2 | 12 | 4.6 (1.8) | 5.0 (4.0-6.0) | 0 | 8 | 4.0 (1.1) | 4.0 (3.0-5.0) | 2 | 6 | 15.8 (1.4) | 16.0 (15.0-17.0) | 14 | 19 | 44.8 |
| 7 | 37 | 6.5 (1.8) | 6.0 (5.0-8.0) | 3 | 11 | 12.5 (1.3) | 12.0 (12.0-14.0) | 10 | 14 | 5.6 (1.4) | 6.0 (5.0-7.0) | 3 | 8 | 19.4 (1.8) | 20.0 (18.0-21.0) | 15 | 22 | 56.8 |
| 8 | 23 | 8.3 (1.6) | 8.0 (7.0-10.0) | 5 | 11 | 4.2 (2.1) | 4.0 (3.0-6.0) | 0 | 8 | 5.9 (1.9) | 6.0 (4.0-8.0) | 3 | 10 | 9.7 (1.1) | 10.0 (9.0-11.0) | 8 | 12 | 60.9 |
| 9 | 46 | 9.0 (1.3) | 9.0 (8.0-10.0) | 5 | 12 | 12.4 (1.3) | 13.0 (11.0-13.0) | 10 | 14 | 9.2 (1.8) | 9.0 (8.0-10.0) | 6 | 12 | 22.7 (1.4) | 22.5 (22.0-24.0) | 20 | 25 | 69.6 |
| 10 | 43 | 6.6 (1.4) | 7.0 (6.0-7.0) | 4 | 9 | 11.7 (1.5) | 12.0 (10.0-13.0) | 9 | 14 | 5.0 (1.5) | 5.0 (4.0-6.0) | 2 | 8 | 24.1 (1.3) | 24.0 (23.0-25.0) | 22 | 26 | 55.8 |
| 11 | 34 | 4.4 (1.6) | 5.0 (3.0-6.0) | 0 | 7 | 4.9 (1.8) | 5.0 (4.0-6.0) | 1 | 9 | 4.1 (1.7) | 4.0 (3.0-5.0) | 1 | 9 | 10.9 (1.9) | 10.5 (9.0-13.0) | 8 | 14 | 55.9 |
| 12 | 38 | 8.8<br>(1.6) | 9.0<br>(8.0-10.0) | 5 | 12 | 9.7<br>(1.5) | 9.0<br>(8.0-11.0) | 8 | 13 | 9.2<br>(1.3) | 9.0<br>(8.0-10.0) | 7 | 12 | 18.1<br>(1.3) | 18.0<br>(17.0-19.0) | 15 | 20 | 65.8 |
| 13 | 52 | 7.1<br>(2.0) | 7.0<br>(5.5-8.5) | 3 | 12 | 7.4<br>(1.5) | 7.0 (6.0-9.0) | 4 | 10 | 4.6<br>(1.5) | 4.5<br>(3.5-6.0) | 1 | 7 | 22.0<br>(1.7) | 22.0<br>(21.0-23.0) | 19 | 27 | 44.2 |
| 14 | 46 | 7.9<br>(1.5) | 8.0<br>(7.0-9.0) | 4 | 11 | 6.2<br>(1.6) | 6.0 (5.0-7.0) | 1 | 9 | 8.4<br>(1.7) | 8.0<br>(7.0-10.0) | 5 | 12 | 19.4<br>(1.4) | 19.0<br>(18.0-20.0) | 17 | 22 | 58.7 |
| 15 | 35 | 9.5<br>(1.4) | 9.0<br>(8.0-11.0) | 7 | 12 | 12.7<br>(1.4) | 13.0<br>(12.0-14.0) | 9 | 14 | 9.6<br>(1.1) | 10.0<br>(9.0-10.0) | 8 | 12 | 27.5<br>(1.5) | 27.0<br>(26.0-29.0) | 25 | 30 | 54.3 |

ADI-R: Autism Diagnostic Interview-Revised.
SD: standard deviation.

### Gene interpretation

We observed 65 chromosomal loci that satisfied the threshold of *P* < 5.0 × 10^−8^ (Table 1 and Figure 3); 30 out of the 65 loci were located within 21 genes, and the remaining 35 loci were intergenic (Table 3). Among them, 8 loci were located within or near the genes associated with the Human Gene module of the SFARI Gene scoring system (6); *GABBR2* (score 4, Rare Single Gene Mutation, Syndromic, Functional) in Cluster 1; *CNTNAP5* (score 4, Rare Single Gene Mutation, Genetic Association) in Cluster 3; *ITPR1* (score 4, Rare Single Gene Mutation) in Cluster 5; *DNAH17* (score 4, Rare Single Gene Mutation) in Cluster 7; *SDK1* (score none, Rare Single Gene Mutation, Genetic Association) in Cluster 13; *SRRM4* (score 5, Rare Single Gene Mutation, Functional) in Cluster 13; *CNTN5* (score 3, Rare Single Gene Mutation, Genetic Association) in Cluster 14; and *DPP10* (score 3, Rare Single Gene Mutation) in Cluster 15.

**Table 3.**
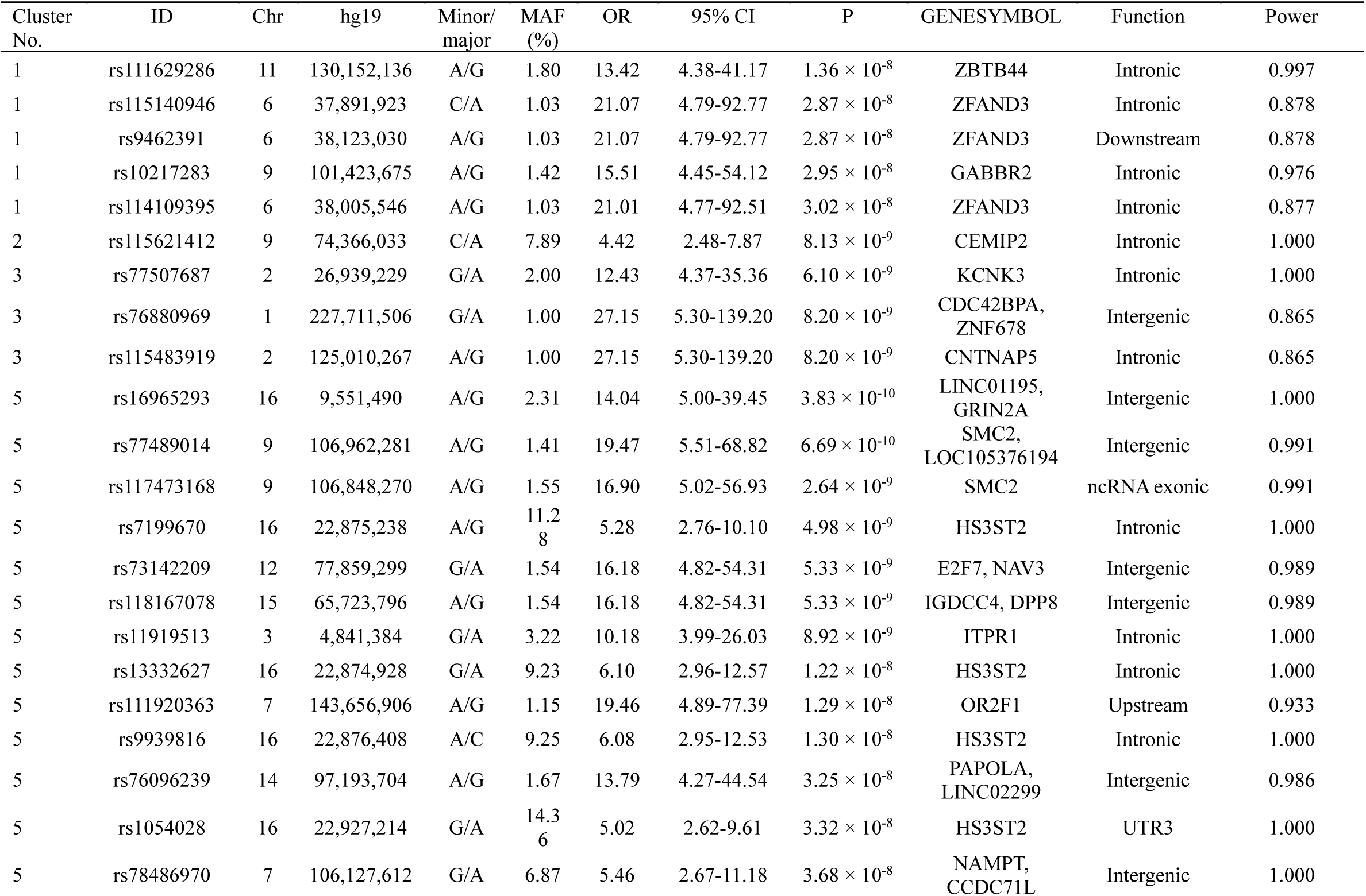
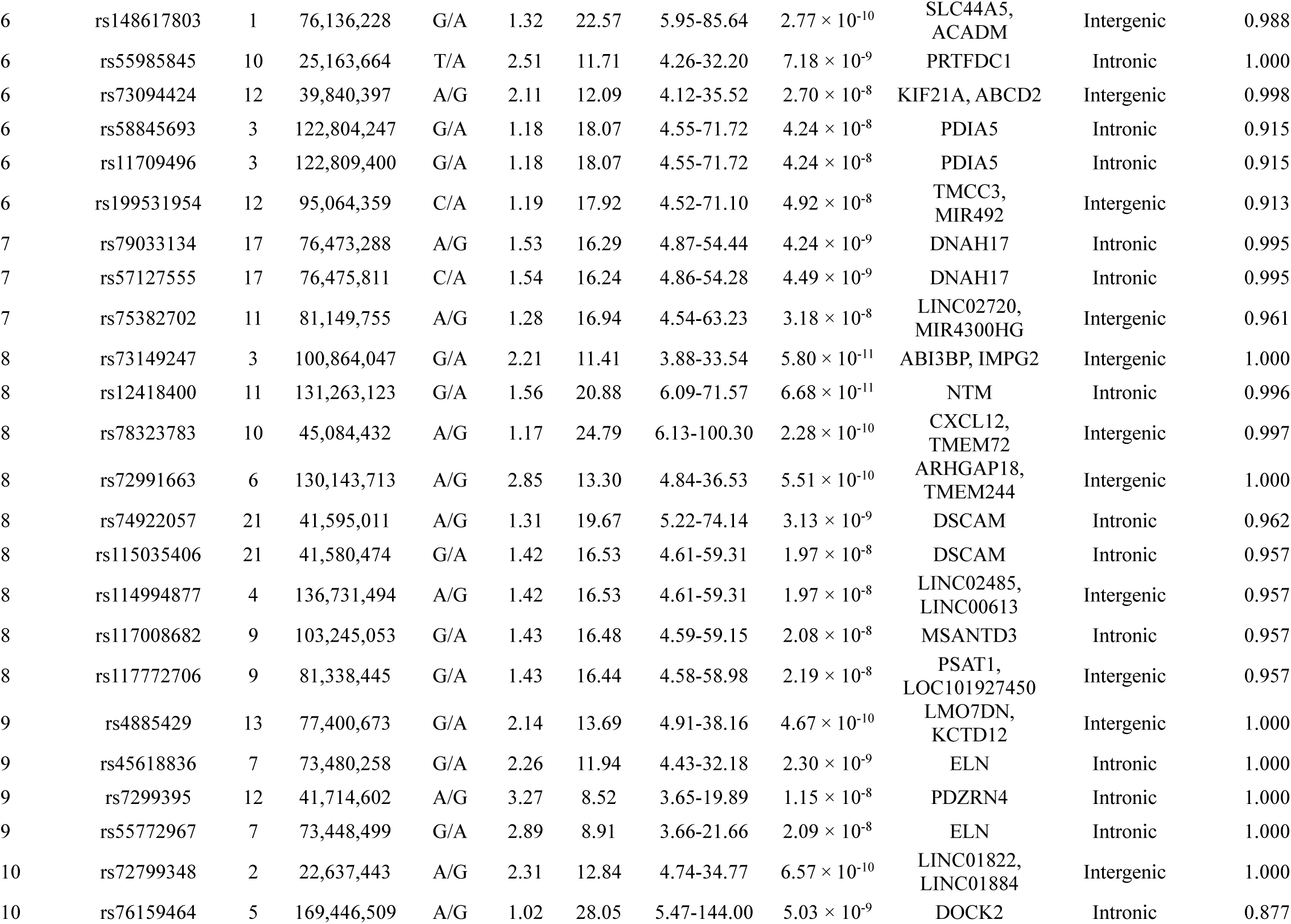
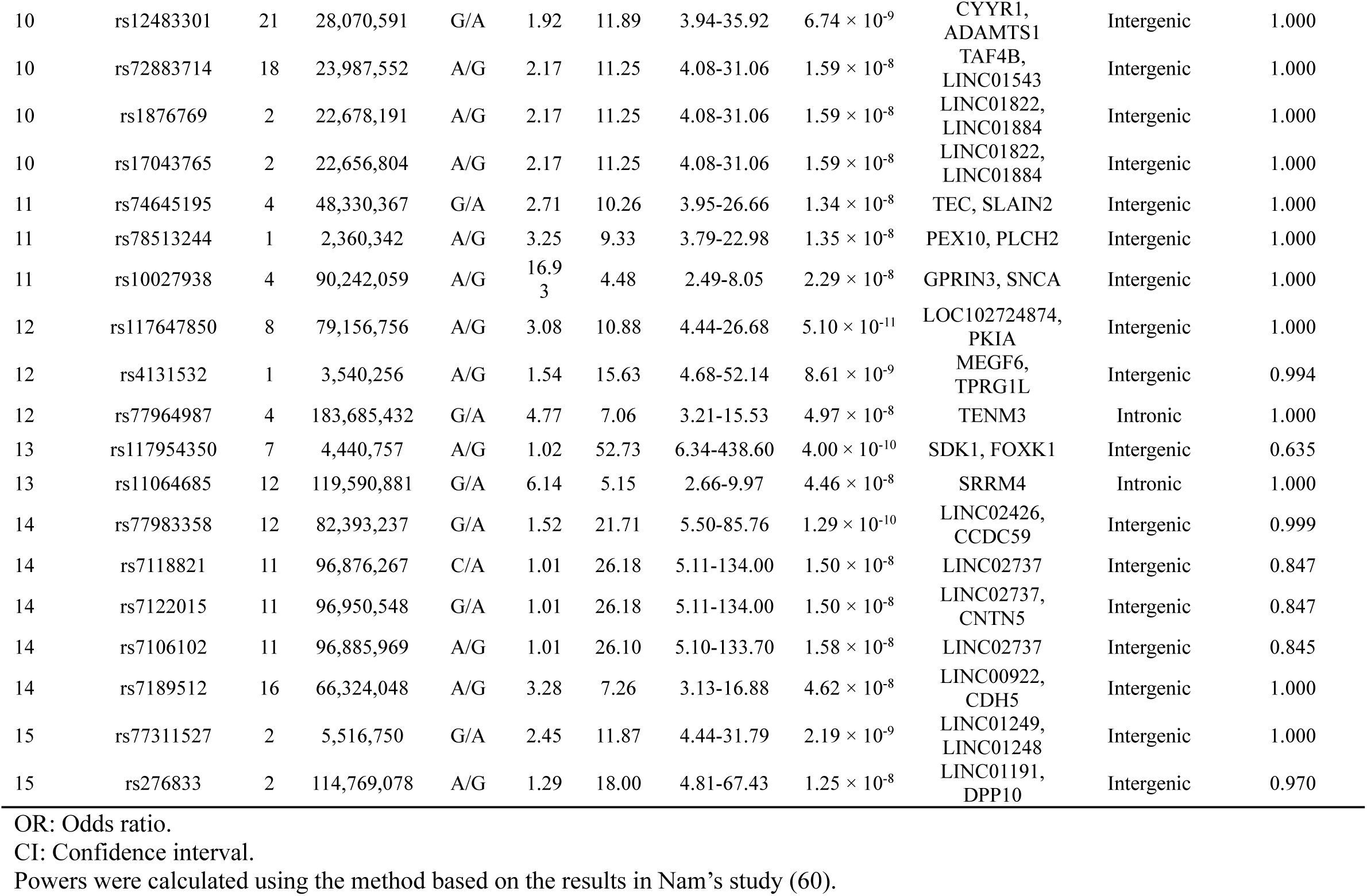
Association table of the cluster-based GWAS with 15 k-means clusters in the Omni2.5 dataset.

The SFARI Gene scoring system ranges from “Category 1”, which indicates “high confidence”, through “Category 6”, which denotes “evidence does not support a role”. Genes of a syndromic disorder (e.g., fragile X syndrome) related to ASD are categorized in a different category. Rare single gene variants, disruptions/mutations, and submicroscopic deletions/duplications related to ASD are categorized as “Rare Single Gene Mutation”.

In addition to genes in the Human Gene module of the SFARI Gene, several important genes associated with ASD or other related disorders (34-37) from previous reports were included in our findings as follows: *CDH5* in Cluster 14, *DSCAM* in Cluster 8, *FOXK1* in Cluster 13, *GRIN2A* in Cluster 5, *NTM* in Cluster 8, and *SNCA* in Cluster 11 previously reported with ASD (38-43); *PLCH2* in Cluster 11 previously reported with mental retardation (44); *ARHGAP18* in Cluster 18, *CDC42BPA* in Cluster 3, *CXCL12* in Cluster 8, and *HS3ST2* in Cluster 5 previously reported with schizophrenia (45-48); *KCTD12* in Cluster 9 and *PSAT1* in Cluster 8 previously reported with depressive disorder (49, 50); *ADAMTS1* in Cluster 10, *DOCK2* in Cluster 10, *HS3ST2* in Cluster 5, *NAMPT* in Cluster 5, and *NAV* in Cluster 5 previously reported with Alzheimer’s disease (51-55); and *PEX10* in Cluster 11 previously reported with Down syndrome (56).

### Replication study

We conducted replication studies with another independent dataset that included a total of 712 male probands and 354 unaffected brothers and had been genotyped using the 1Mv3 array. As mentioned before, we had previously carried out cluster analyses in the combined dataset genotyped with either Omni2.5 or 1Mv3 and then redivided it according to the SNP arrays used. The characteristics of each of the 15 clusters in the 1Mv3 dataset are presented in Table S1.

Among the 65 genome-wide significant chromosomal loci found in the discovery study, seven chromosomal loci were included in the 1Mv3 array. Of these loci, rs11064685, within *SRRM4* in Cluster 13, had a significantly different distribution (p =0.03) in cases vs. controls in the replication cohort (Table 4).

**Table 4.** Results of replication studies in the 1Mv3 dataset for statistically significant chromosomal loci in the discovery studies.

| Cluster No. | ID | Chr | hg19 | Minor/<br>major | MAF (%) | OR | 95% CI | P | GENESYMBOL | Function |
| --- | --- | --- | --- | --- | --- | --- | --- | --- | --- | --- |
| 5 | rs13332627 | 16 | 22,874,928 | G/A | 10.0 | 0.50 | 0.18-1.45 | 0.195 | HS3ST2 | intronic |
| 5 | rs7199670 | 16 | 22,875,238 | A/G | 12.2 | 0.51 | 0.20-1.33 | 0.1629 | HS3ST2 | intronic |
| 5 | rs1054028 | 16 | 22,927,214 | G/A | 15.0 | 0.51 | 0.22-1.21 | 0.121 | HS3ST2 | UTR3 |
| 10 | rs1876769 | 2 | 22,678,191 | A/G | 1.4 | NA | - | 0.1822 | LINC01822,<br>LINC01884 | intergenic |
| 13 | rs11064685 | 12 | 119,590,881 | G/A | 8.2 | 1.89 | 1.06-3.37 | 0.02858 | SRRM4 | intronic |
| 14 | rs7189512 | 16 | 66,324,048 | A/G | 3.5 | 2.16 | 0.83-5.67 | 0.1085 | LINC00922,<br>CDH5 | intergenic |
| 15 | rs276833 | 2 | 114,769,078 | A/G | 1.3 | 0.71 | 0.09-5.75 | 0.75 | LINC01191,<br>DPP10 | intergenic |

OR: Odds ratio.
CI: Confidence interval.

## Discussion

One of the most important findings of our study was that reasonably decreasing the sample size could increase the statistical power. A plausible explanation is that our clustering may have successfully identified subgroups that are etiologically more homogeneous. At least three reasons could reduce the possibility of false positives of the present results of statistically significant SNPs in cluster-based GWAS. First, the present study validated the usefulness and feasibility of the concept of a previous simulation study (8), which indicated that homogeneous case subgroups increase power in genetic association studies by Traylor and colleagues, using measurement data in the real world. Second, a substantial number of statistically significant SNPs in cluster-based GWAS observed in the present study were located within or near previously reported candidate genes for ASD (6, 38-43). Third, we calculated λ of Q-Q plots of logarithm of P-values to base 10 (-log10P) for each cluster number and compared them for the validity of clustering. Genomic control is widely used to control false positive signals due to population stratification in GWAS. λ is commonly used for genomic control.

We observed many statistically significant SNPs in cluster-based GWAS: *CDH5, CNTN5, CNTNAP5, DNAH17, DPP10, DSCAM, FOXK1, GABBR2, GRIN2A*5, *ITPR1, NTM, SDK1, SNCA* and *SRRM4.* In particular, loci within the *SRRM4* gene had significantly different distributions in the cases vs. controls in the replication cohort. Previous studies indicate that *SRRM4* is strongly associated with ASD, indicating that our results may be valid to some degree. The gene regulates neural microexons. In the brains of individuals with ASD, these microexons are frequently dysregulated (57). Additionally, nSR100/SRRM4 haploinsufficiency in mice induced autistic features such as sensory hypersensitivity and altered social behavior and impaired synaptic transmission and excitability (58).

In addition to *SMMR4*, we observed several genes located within or near previously reported candidate genes for ASD. The relatively high correspondence between our results in part and the SFARI Gene scoring system (6) indicates that the statistically significant loci we found may be associated with ASD subgroups (Table 1 and Figure 3). We also observed several important genes associated with ASD and other related disorders (34-37) from previous reports. These findings suggest that the statistically significant SNPs might explain autistic symptoms because these diseases are suggested to have shared etiology, even in part, with ASD (34-37). Associations at the remaining significant loci that were not in the SFARI module or described above have not been previously reported, and to the best of our knowledge, some of them might be novel findings. These results might suggest that novel genetic loci of ASD could be found by identifying better defined subgroups, although further confirmation is needed in future cohorts with larger sample sizes.

Previous studies regarding Alzheimer’s disease, neuroticism, or asthma found that items or symptoms showed, to some degree, increased ORs between the case loci and control loci compared to those from previous studies using broadly defined disease diagnoses (9-11). These findings may indicate that GWAS based on a symptom or an item could identify genetically more homogeneous subgroups and let us hypothesize that a relatively reasonable combination of symptoms or items could identify more genetically homogeneous subgroups.

In contrast, Chaste and colleagues showed that stratifying children with ASD based on the phenotype only modestly increased power in GWAS (12). The discrepancy between their findings and ours might be explained by at least two reasons. First, Chaste and colleagues used one item or symptom alone, whereas we used combinations of them with a machine learning method. DeMichele-Sweet and colleagues reported that subgrouping only by having psychosis could lead to the identification of limited loci that had small effects (59), but Mukherjee and colleagues found a substantial number of suggestive loci that had extreme ORs after categorizing persons with Alzheimer’s disease based on relative performance across cognitive domains by modern psychometric approaches (9). It may be necessary to utilize an appropriate combination of data to reveal masked patterns of data sets. Second, the number of subgroups was quite different between Chaste and colleagues’ study and ours. Chaste and colleagues divided their participants mainly into two subgroups, but we divided ours into 15. In fact, when we employed a cluster number of two in our study, we observed no significant loci (Table 1). If ASD consists of more than hundreds of subgroups (6), grouping with a sufficient number of clusters may be necessary.

Validation of clusters is essential. In the present study, we selected the k-means algorithm, focused on ADI-R items and treatment as variables, and determined cluster numbers based on the λ of the Q-Q plots. Although we believe this approach is one of the relevant ways, selection of variables, selection of algorithms and selection of cluster numbers still remain to be considered in future mathematical and biological cluster validation studies because controversies surrounding evaluation of the quality of the clusters are important issues and are still ongoing and because validated clusters may lead to elucidate the genetic architectures of ASD (24).

The present study has a limitation to be noted. Substantial differences in the two genotyping platforms may have affected the results of the replication study. The Omni2.5 array includes 2,383,385 autosomal SNPs, whereas the 1Mv3 array includes 1,147,689 SNPs, with 675,923 shared SNPs between the two. Of the 65 statistically significant chromosomal loci in the discovery data, only seven chromosomal loci were shared between the two arrays.

Our study demonstrated that if the data set consists of multiple heterogeneous subgroups, even a subgroup that includes a much smaller number of homogeneous individuals could detect high-impact genetic factors. Hypothetical examples of the concept of cluster-based GWAS are shown in Figure S3. As shown in this figure, in the conventional design in which a whole data set is involved, an actual effect of a variant would be “diluted” to a modest OR of, e.g., 1.5, and at least thousands or tens of thousands of individuals would be required to detect it as a significantly associated variant. In contrast, cluster-based GWAS would be more likely than the conventional design to detect associated variants, without their effects being diluted, and with much higher ORs. As shown in Table 3, only 30 etiologically homogeneous probands and 300 controls can have a statistical power of approximately 1.00, calculated using the method based on the results in Nam’s study (60). Although the integral model, which assumes many genetic variants have a small effect, may contribute to the formation of some subgroups of ASD, our results indicate that clustering by specific phenotypic variables may provide a candidate example for identifying etiologically similar cases of ASD.

Our data indicate the relevance of cluster-based GWAS as a means to identify more homogeneous subgroups of ASD than broadly defined subgroups. Future investigation of cluster validation and replication with a larger sample size is therefore warranted. Such studies will provide clues to elucidate the genetic structures and etiologies of ASD and facilitate the development of precision medicine for ASD.

## Acknowledgments

We are grateful to all of the families at the participating Simons Simplex Collection (SSC) sites, as well as the staff at the Simons Foundation Autism Research Initiative (SFARI). The present study was supported by the Ministry of Education, Culture, Sports, Science and Technology (MEXT) KAKENHI Grant Numbers 19390171 and 16H05242. MEXT had no role in the design or execution of the study.

