## Supplementary Table S1 for "Clustering by phenotype and genome-wide association study in autism"

**Table S1.** The characteristics of each cluster divided by 15 clusters in the 1Mv3 dataset.

| Cluster No. | Verbal score from ADI-R | | | | Non-verbal score from ADI-R | | | | Restricted and repetitive patterns of behavior scor from ADI-R | | | |
|---|---|---|---|---|---|---|---|---|---|---|---|---|
| | Mean (SD) | Median (p25-p75) | Min | Max | Mean (SD) | Median (p25-p75) | Min | Max | Mean (SD) | Median (p25-p75) | Min | Max |
| All (n=712) | 7.5 (2.0) | 8.0 (6.0-9.0) | 1 | 12 | 8.7 (3.4) | 9.0 (6.0-12.0) | 1 | 14 | 6.6 (2.6) | 6.0 (5.0-8.0) | 0 | 12 |
| 1 (n=44) | 7.8 (1.9) | 8.0 (6.0-9.0) | 5 | 12 | 4.4 (1.6) | 5.0 (3.0-6.0) | 1 | 7 | 8.3 (1.4) | 8.0 (7.5-9.0) | 6 | 11 |
| 2 (n=56) | 8.0 (1.7) | 8.0 (7.0-9.0) | 4 | 12 | 12.5 (1.4) | 13.0 (12.0-14.0) | 9 | 14 | 6.5 (1.3) | 7.0 (6.0-7.0) | 3 | 8 |
| 3 (n=57) | 6.2 (1.5) | 6.0 (5.0-7.0) | 3 | 9 | 8.6 (1.3) | 9.0 (8.0-9.0) | 6 | 12 | 4.8 (1.6) | 5.0 (4.0-6.0) | 0 | 8 |
| 4 (n=39) | 8.8 (1.3) | 9.0 (8.0-9.0) | 7 | 12 | 8.1 (1.5) | 8.0 (7.0-9.0) | 5 | 10 | 8.8 (1.6) | 8.0 (8.0-10.0) | 6 | 12 |
| 5 (n=35) | 7.7 (1.4) | 8.0 (7.0-9.0) | 4 | 11 | 9.9 (1.9) | 10.0 (8.0-12.0) | 7 | 14 | 6.3 (2.4) | 6.0 (5.0-8.0) | 2 | 12 |
| 6 (n=47) | 6.9 (1.8) | 7.0 (5.0-8.0) | 4 | 10 | 4.7 (1.5) | 5.0 (4.0-6.0) | 1 | 7 | 4.4 (1.5) | 5.0 (3.0-6.0) | 0 | 7 |
| 7 (n=31) | 6.3 (2.0) | 6.0 (5.0-8.0) | 2 | 11 | 12.5 (1.2) | 12.0 (11.0-14.0) | 10 | 14 | 5.4 (1.9) | 6.0 (4.0-7.0) | 1 | 8 |
| 8 (n=51) | 7.7 (1.5) | 7.0 (7.0-9.0) | 5 | 11 | 4.5 (1.6) | 5.0 (3.0-6.0) | 1 | 8 | 6.1 (1.9) | 6.0 (5.0-7.0) | 3 | 10 |
| 9 (n=44) | 8.8 (1.5) | 9.0 (8.0-10.0) | 5 | 12 | 12.6 (1.1) | 13.0 (12.0-13.0) | 10 | 14 | 8.8 (1.7) | 9.0 (8.0-10.0) | 6 | 12 |
| 10 (n=52) | 6.3 (1.5) | 6.0 (5.0-7.0) | 3 | 9 | 11.7 (1.7) | 12.0 (10.0-13.0) | 7 | 14 | 4.6 (1.9) | 5.0 (3.0-6.0) | 0 | 8 |
| 11 (n=46) | 4.6 (1.5) | 5.0 (4.0-6.0) | 1 | 7 | 4.9 (1.8) | 5.0 (4.0-6.0) | 1 | 10 | 4.2 (1.8) | 4.0 (3.0-5.0) | 1 | 9 |
| 12 (n=46) | 8.2 (1.6) | 8.0 (7.0-9.0) | 5 | 12 | 10.3 (1.6) | 10.0 (9.0-11.0) | 7 | 13 | 9.2 (1.5) | 9.0 (8.0-10.0) | 6 | 12 |
| 13 (n=65) | 7.4 (1.8) | 8.0 (6.0-9.0) | 3 | 11 | 7.6 (1.5) | 8.0 (7.0-9.0) | 3 | 10 | 4.3 (1.3) | 4.0 (3.0-5.0) | 1 | 6 |
| 14 (n=48) | 7.9 (1.8) | 8.0 (6.0-10.0) | 4 | 11 | 6.2 (1.5) | 6.0 (5.0-7.0) | 3 | 8 | 8.5 (1.4) | 8.0 (7.5-9.0) | 6 | 12 |
| 15 (n=51) | 9.3 (1.8) | 10.0 (8.0-11.0) | 6 | 12 | 12.4 (1.4) | 12.0 (11.0-14.0) | 9 | 14 | 9.8 (1.2) | 10.0 (9.0-10.0) | 8 | 12 |

| Cluster No. | Social score from ADI-R | | | | Treatment with vitamin B6 |
|---|---|---|---|---|---|
| | Mean (SD) | Median (p25-p75) | Min | Max | % |
| All (n=712) | 19.1 (5.7) | 19.0 (15.0-24.0) | 8 | 30 | 56.3 |
| 1 (n=44) | 14.2 (1.4) | 14.0 (13.0-15.0) | 12 | 17 | 65.9 |
| 2 (n=56) | 27.0 (1.2) | 27.0 (26.0-27.0) | 25 | 30 | 75.0 |
| 3 (n=57) | 16.6 (1.4) | 17.0 (15.0-18.0) | 14 | 19 | 52.6 |
| 4 (n=39) | 24.3 (1.5) | 24.0 (23.0-26.0) | 22 | 27 | 46.2 |
| 5 (n=35) | 12.8 (1.7) | 13.0 (11.0-14.0) | 9 | 15 | 51.4 |
| 6 (n=47) | 15.4 (1.6) | 15.0 (14.0-17.0) | 13 | 19 | 48.9 |
| 7 (n=31) | 19.8 (1.3) | 20.0 (19.0-21.0) | 17 | 22 | 61.3 |
| 8 (n=51) | 9.5 (1.3) | 9.0 (8.0-10.0) | 8 | 12 | 51.0 |
| 9 (n=44) | 23.0 (1.4) | 23.0 (22.0-24.0) | 20 | 25 | 65.9 |
| 10 (n=52) | 24.2 (1.4) | 24.0 (23.0-25.0) | 22 | 28 | 48.1 |
| 11 (n=46) | 10.7 (1.7) | 11.0 (10.0-12.0) | 8 | 14 | 54.4 |
| 12 (n=46) | 18.2 (1.4) | 18.0 (17.0-19.0) | 16 | 21 | 54.4 |
| 13 (n=65) | 21.2 (1.5) | 21.0 (20.0-22.0) | 19 | 25 | 58.5 |
| 14 (n=48) | 19.2 (1.4) | 19.0 (18.0-20.0) | 17 | 22 | 56.3 |
| 15 (n=51) | 27.6 (1.4) | 27.0 (26.0-29.0) | 26 | 30 | 52.9 |

ADI-R: Autism Diagnostic Interview-Revised.

SD: standard deviation.
