## Supplementary figures and images for "Clustering by phenotype and genome-wide association study in autism"

### Supplementary Figure S1

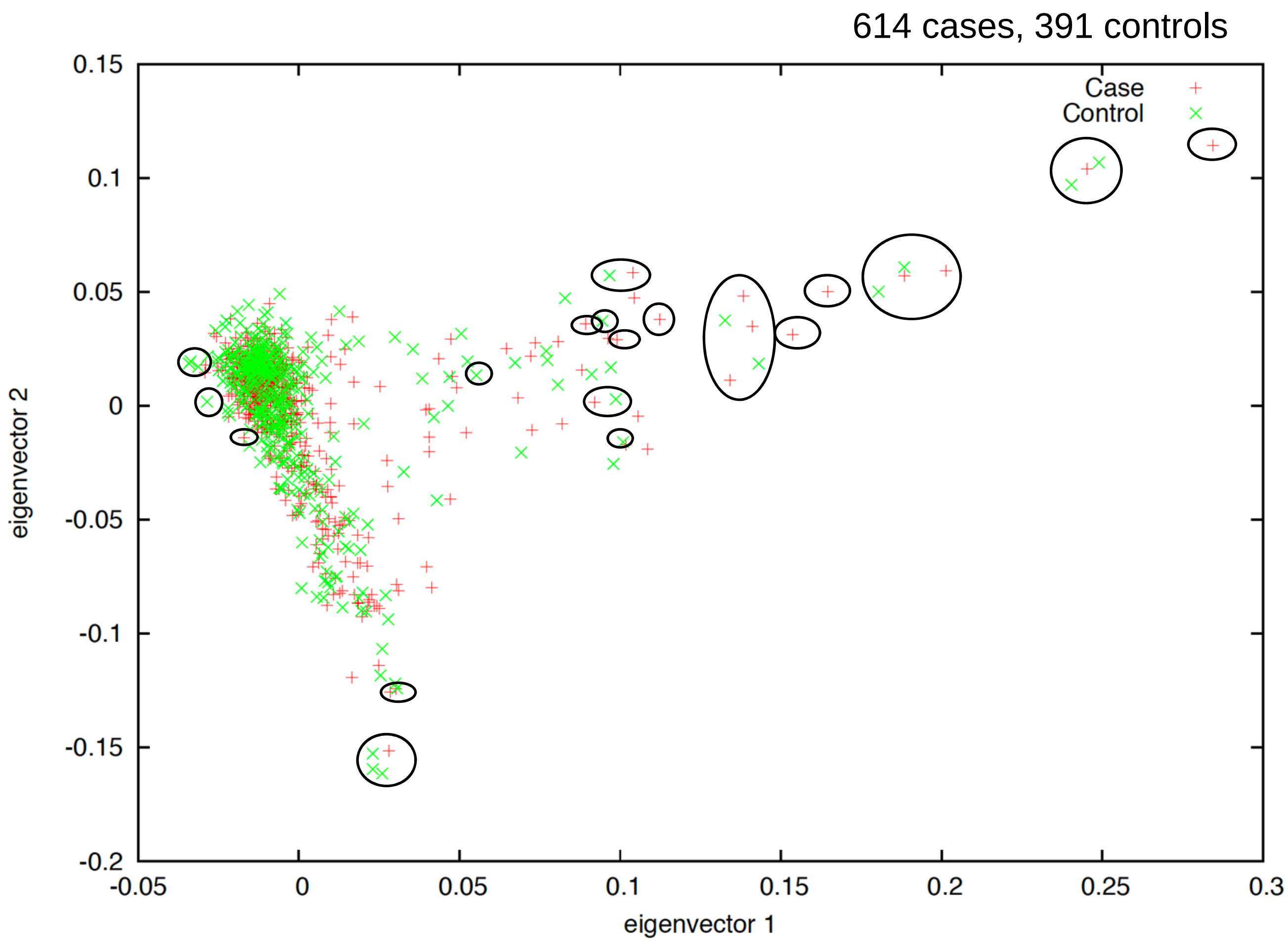


**Figure S1.** Principal component analysis of the genotyping data in the discovery stage.
