## Supplementary Figure S2 for "Clustering by phenotype and genome-wide association study in autism"

Cochran–Armitage trend test for the whole dataset

Sib-TDT test for one subset of probands of cluster 1 vs all the controls using k-means with 15 clusters

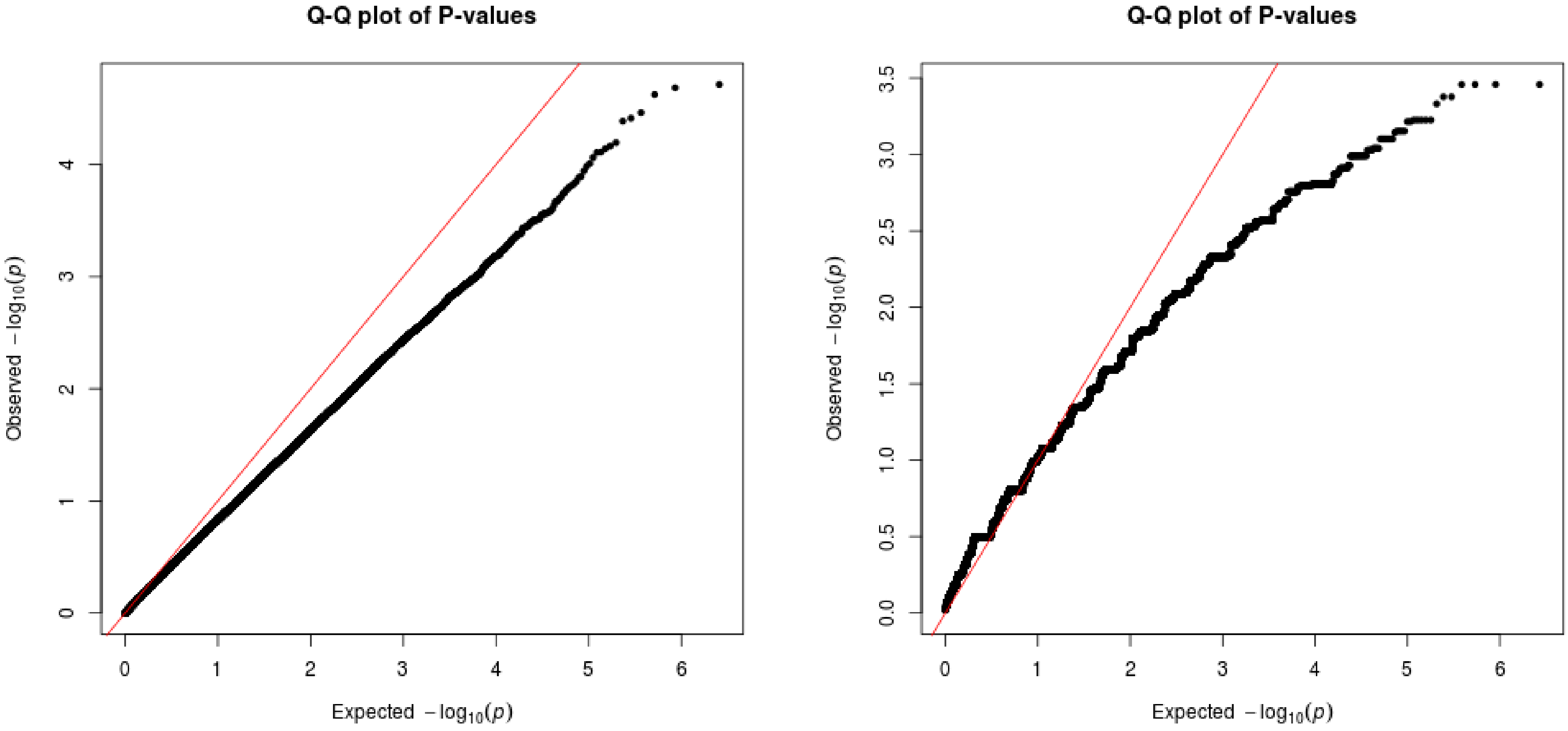


**Figure S2.** The Cochran–Armitage trend test for the whole dataset, and the sib-TDT test for one subset of probands of cluster 1 vs all the controls.
