## Supplementary Figure S3 for "Clustering by phenotype and genome-wide association study in autism"

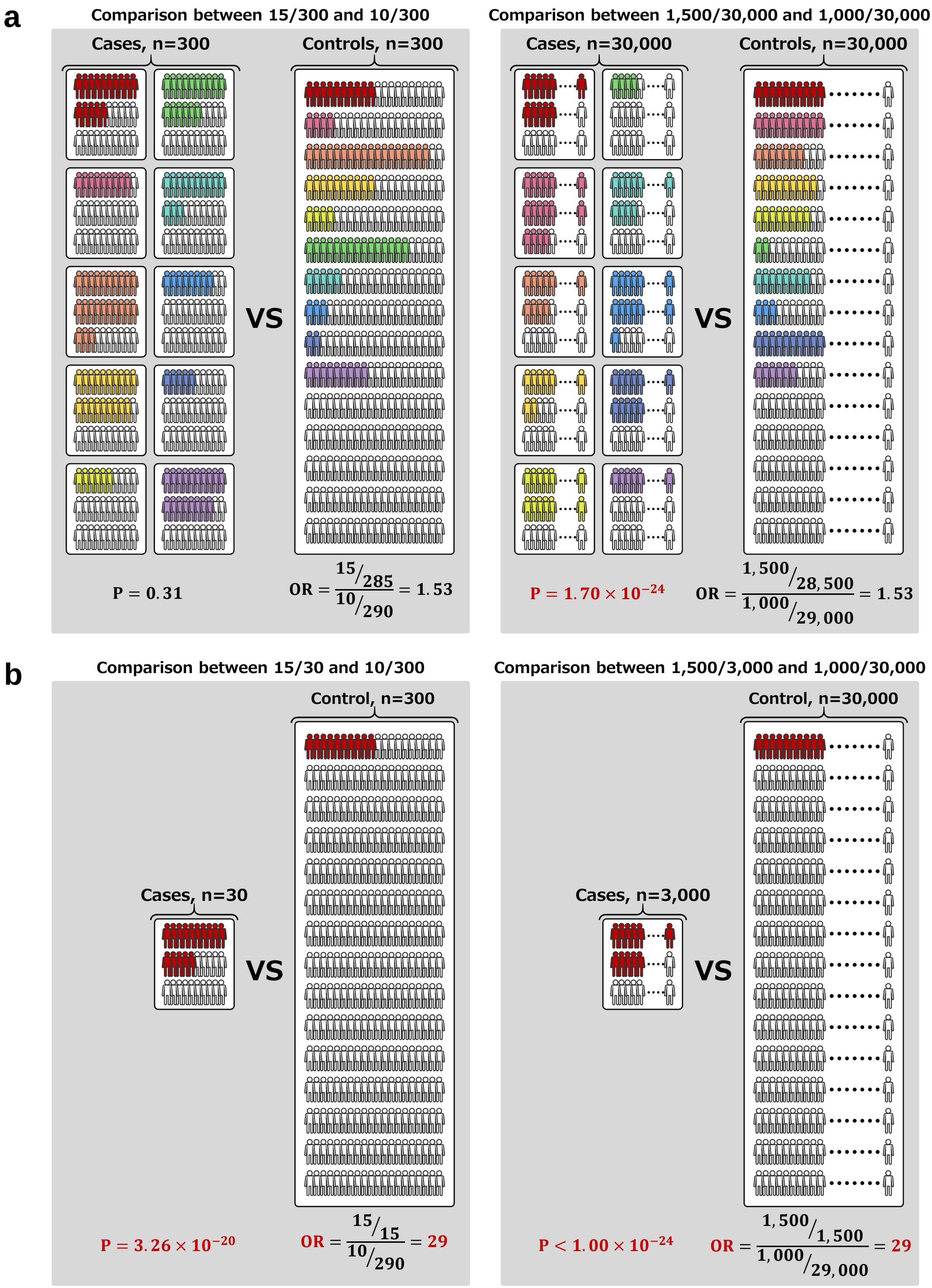


**Figure S3.** Hypothetical examples of the concept of cluster-based GWAS. GWAS for all cases vs all controls are shown in (**a**) and cluster-based GWAS are shown in (**b**). These examples are based on the hypothesis that autism spectrum disorders might consist of ten causes. In this example of persons described in red colour, in the all cases vs all controls analysis, analysis with a large sample size revealed significant loci, although the odds ration (OR) remained low. In contrast, in the cluster-based GWAS, both analyses revealed significant loci and a large OR.
