## Supplementary Information S1 for "Clustering by phenotype and genome-wide association study in autism"

**Supplementary Information S1.** Python code of kmeans

```
import pandas as pd
f=pd.read_csv("ssc.csv")
import matplotlib.pyplot as plt
from sklearn.cluster import KMeans
model1=KMeans(n_clusters=2, random_state=0)
model2=KMeans(n_clusters=3, random_state=0)
model3=KMeans(n_clusters=4, random_state=0)
model4=KMeans(n_clusters=5, random_state=0)
model5=KMeans(n_clusters=10, random_state=0)
model6=KMeans(n_clusters=15, random_state=0)
model7=KMeans(n_clusters=20, random_state=0)
data=f[['adi_r_comm_only_verbal_total','adi_r_comm_b_non_verbal_total','adi_r_rrb_c_tota
l','adi_r_soc_a_total','child_vitamins']]
model1.fit(data)
model2.fit(data)
model3.fit(data)
model4.fit(data)
model5.fit(data)
model6.fit(data)
model7.fit(data)
y=model1.labels_
y2=model2.labels_
y3=model3.labels_
y4=model4.labels_
y5=model5.labels_
y6=model6.labels_
y7=model7.labels_
data1=f.copy()
data2=f.copy()
data3=f.copy()
data4=f.copy()
data5=f.copy()
data6=f.copy()
data7=f.copy()
```

```
data1['cluster']=y
data2['cluster']=y2
data3['cluster']=y3
data4['cluster']=y4
data5['cluster']=y5
data6['cluster']=y6
data7['cluster']=y7
data1.to_csv("kmeans(c=2).csv")
data2.to_csv("kmeans(c=3).csv")
data3.to_csv("kmeans(c=4).csv")
data4.to_csv("kmeans(c=5).csv")
data5.to_csv("kmeans(c=10).csv")
data6.to_csv("kmeans(c=15).csv")
data7.to_csv("kmeans(c=20).csv")
```
